# Unilateral vocal nerve resection alters neurogenesis in the avian song system in a region-specific manner

**DOI:** 10.1101/2020.03.24.005330

**Authors:** Jake V. Aronowitz, Alice Perez, Christopher O’Brien, Siaresh Aziz, Erica Rodriguez, Kobi Wasner, Sissi Ribeiro, Dovounnae Green, Farhana Faruk, Carolyn L. Pytte

**Author notes:** Equal contribution (Jake V. Aronowitz, Alice Perez).

## Abstract

New neurons undergo a critical period soon after migration during which the behavior of the animal may result in the survival or culling of these cells. In the songbird song system, new neurons may be maintained in the song motor pathway with respect to motor progression toward a target song--during juvenile song learning, seasonal song restructuring, and experimentally manipulated song variability. However, it is not known whether the quality of song per se, without progressive improvement, may also influence new neuron survival. To test this idea, we experimentally altered song acoustic structure by unilateral denervation of the syrinx. We found no effect of aberrant song on numbers of new neurons in the HVC of the song motor pathway, a loss of left-side dominance in new neurons in the auditory region caudomedial nidopallium (NCM), and a bilateral decrease in new neurons in the basal ganglia nucleus Area X. We propose new neuron survival may be determined in response to behavioral feedback in accordance with the function of new neurons within a given brain region. Studying the effects of singing behaviors on new neurons across multiple brain regions that subserve singing may give rise to general rules underlying the regulation of new neuron survival across taxa and brain regions more broadly.

## Introduction

Most cell types that continue to be produced postnatally have predetermined lifespans, varying by tissue type, and largely independent of the experience of the cell. In healthy adults, epithelial cells survive less than 2 weeks (Spalding et al., 2005), white blood cells live for about 13-20 days, red blood cells survive 100-120 days (Krzyzanski et al., 2013), and liver hepatocytes turnover about every 200-300 days (Macdonald, 1961; Stocker and Heine, 1971). On the other hand, across species and brain regions, lifespans of neurons born in the postnatal brain are highly variable. More interestingly, lifespans differ not only by neuron type, but are influenced by the cell’s activity and experience, setting them apart in this way from other continually regenerating cell types (Kirn et al., 1994; Kirn and Schwabl, 1997; Gheusi et al., 2000; Rochefort et al., 2002; Shors et al., 2002; Dupret et al., 2008; Belnoue et al., 2013; Liu and Nusslock, 2018; Larson et al., 2019).

Time-dependent changes in the expression of a number of transmembrane factors and intracellular signaling molecules in immature neurons seem to mediate the transition between cell survival and cell death, with different mosaics of receptors expressed throughout neuronal maturation in a cell-type specific manner (Pfisterer and Khodosevich, 2017). However, even after an early critical period of culling within the first few weeks of a neuron’s life, microenvironmental and behavioral conditions continue to impact the neuronal lifespan (Kirn et al., 1999; Alvarez-Borda et al., 2004; Brenowitz and Larson, 2015; Shevchouk et al., 2017).

In the songbird, behavioral factors affecting new neuron survival have been studied most systematically in the nucleus HVC (Wilbrecht and Kirn, 2004; Brenowitz and Larson, 2015), a sensorimotor region subserving song production that also responds to song auditory stimuli (McCasland, 1987; George et al., 2005). Throughout the bird’s life, HVC receives new neurons that project to the premotor robust nucleus of the arcopallium (RA), becoming part of the motor pathway for song production (HVC_RA_). In the canary, amount of singing positively correlates with new neuron survival (Li et al., 2000; Alvarez-Borda and Nottebohm, 2002; Ball et al., 2004). This association is mediated by BDNF and testosterone, both of which may directly increase new neuron survival (Rasika et al., 1994; Rasika et al., 1999; Alvarez-Borda et al., 2004; Brenowitz and Larson, 2015; Alward et al., 2016; Ball et al., 2020). Housing conditions also impact the lifespan of both HVC _RA_ neurons and new neurons in the auditory region caudomedial nidopallium (NCM), with more new neurons seen in zebra finches or canaries housed in groups compared to singly housed birds (Lipkind et al., 2002; Barnea et al., 2006; Walton et al., 2012; Shevchouk et al., 2017). Interestingly, Shevchouk et al., (2017) show that new neuron proliferation and the subsequent survival of new neurons can be independently affected by male- or female-paired housing. Numerous behaviors may underlie this effect, including increased singing and increased exposure to songs. Intriguingly, environmental factors of photo period and temperature appear to influence new neuron numbers in the caudal nidopallium and hippocampus in the non-seasonal zebra finch (Pozner et al., 2018).

Across taxa, a common modulator of neuronal lifespan is neural activity, with most evidence indicating that activity confers survival (Alvarez-Borda and Nottebohm, 2002; Ninkovic et al., 2007; Larson et al., 2013; Brenowitz and Larson, 2015; Pfisterer and Khodosevich, 2017; Hall and Tropepe, 2018). However, neuronal activity has also decreased new neuron survival of olfactory periglomerular cells under precise conditions, perhaps fine-tuning olfactory sensitivity to odorant density (Khodosevich et al., 2013). During brain development, neuronal activity conferring cell survival may direct pathway maintenance and pruning thereby building system efficiency. Perhaps a similar functional outcome occurs in adulthood. This has been proposed by Wilbrecht et al. (2002) and (Wilbrecht and Kirn, 2004) in a model in which new HVC neurons audition for a spot in the song production pathway and get the part when their contribution improves song performance. Interestingly, this model considers not only the amount of songs produced, but also the ongoing *quality* of the song. In this way, they propose that the act of accurate progression in a goal directed behavior interacts with song production in promoting new neuron retention (Wilbrecht and Kirn, 2004; Pytte et al., 2008).

Consistent with this idea, experimentally altered song structure, by reversible paralysis of the syringeal muscles, resulted in a positive correlation between numbers of new neurons in HVC and song recovery (Pytte et al., 2011). However, the direction of causality could not be determined. Increasingly accurate song production during song recovery may have promoted the survival of new neurons or birds with higher numbers of new neurons may have been better able to correct altered song structure (Pytte et al., 2011). Song recovery, i.e., goal directed improvement in the behavior, was also confounded by song quality per se. Songs that were the most aberrant recovered the least. Therefore, although consistent with the model that behavioral improvement in quality promotes new neuron survival, either song quality or song recovery may have influenced new neuron survival in this study. Alternatively, perhaps new neurons in HVC drove song recovery and song feedback was not a factor in neuronal survival.

To address this, we tested whether song quality without recovery impacts new neuron survival. Here we produced an irreversibly aberrant song by unilaterally denervating the syrinx in adult male zebra finches, resulting in stable, altered feedback during song production. This allowed us to determine directly whether song quality may impact numbers of new neurons, and also to broaden our investigation to song regions outside of HVC.

In addition to HVC and NCM, the basal ganglia nucleus Area X processes song-related feedback and continues to receive new neurons throughout adulthood (Kosubek-Langer et al., 2017). Area X receives projections from a different population of HVC neurons (HVC_X_) than those that project to RA and is part of the anterior forebrain pathway necessary for juvenile song learning (Sohrabji et al., 1990) and adult song plasticity (Woolley et al., 2014). NCM affects HVC activity indirectly (Pawlisch and Remage-Healey, 2015) and perhaps is a source of auditory input into the “song system,” the group of interconnected nuclei that subserve song behavior (Figure 1). Given the functional and anatomical connections between HVC, NCM and Area X, we explored effects of altered song feedback on new neurons across these regions.

**Figure 1.**
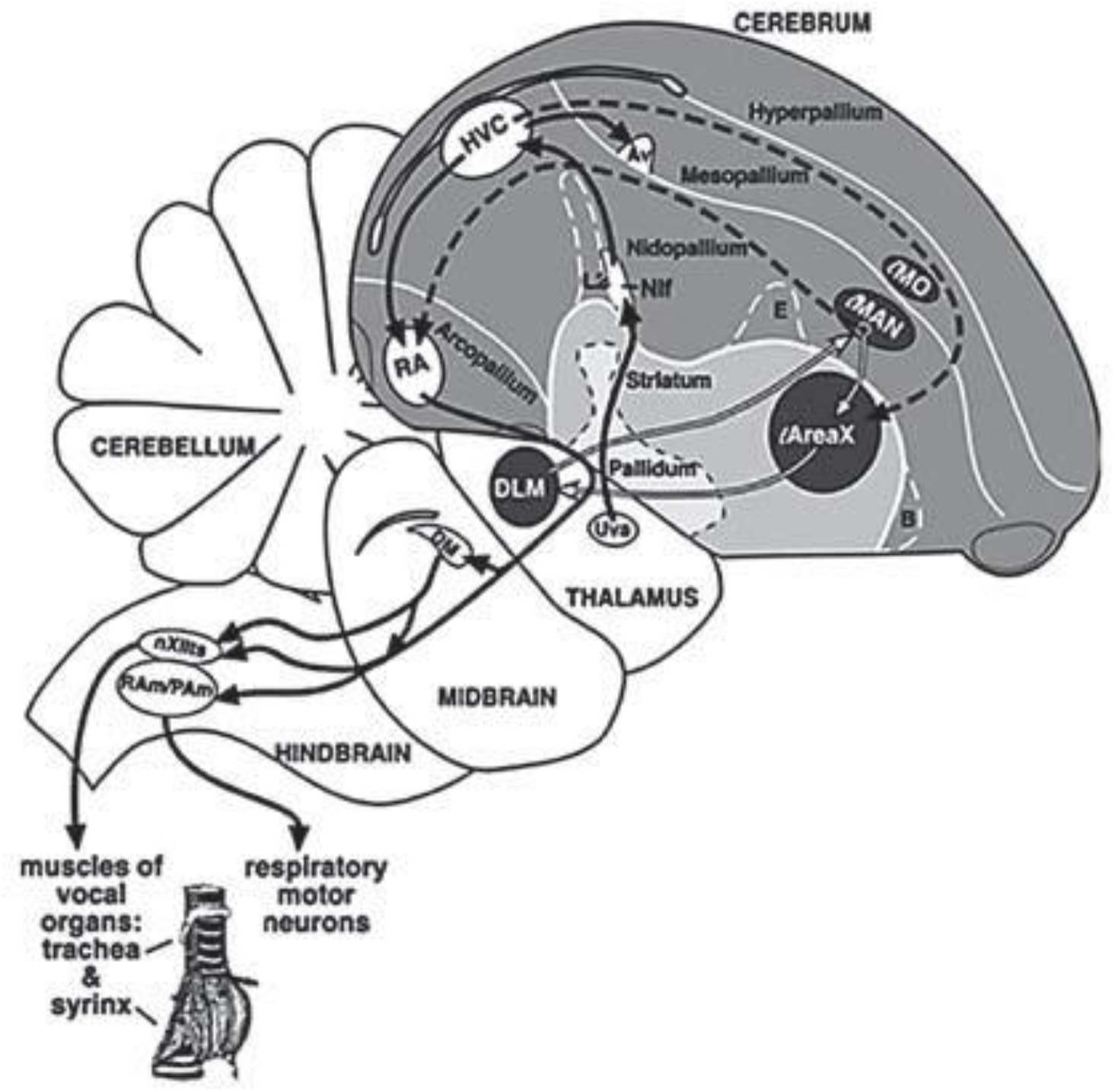
Diagram of the song system in a sagittal section, showing the vocal motor pathway (black solid lines) and the anterior forebrain pathway (dashed and white lines). White lines show the feedback loop between the striatum, thalamus, nidopallium, and projection back to the striatum. AV = Nucleus avalanche; B = nucleus basorostralis; DM = dorsal medial nucleus; DLM = dorsal lateral nucleus of the medial thalamus; E = entopallium; LMAN = lateral magnocellular nucleus of the anterior nidopallium; LMO = lateral oval nucleus of the mesopallium; Nif = interfacial nucleus of the nidopallium; PAm = para-ambiguus; Ram = nucleus retroambiguus; Uva, nucleus uvaeformis; nXIIts = tracheosyringeal portion of the hypoglossal nucleus. Reproduced from Pytte et al., (2011); originally printed in Wilbrecht and Kirn (2004). Syrinx drawn by Franz Goller.

We also speculated that differences in the effects of feedback on new neurons may not only occur across brain regions, but also perhaps across the left and right hemispheres of a given brain region. Tsoi et al. (2014) reported inter-hemispheric differences whereby a greater number of new neurons are added to left NCM than right NCM of adult zebra finches. Moreover, the degree of asymmetry was positively correlated with the quality of song learning and the strength of neuronal memory for songs (Tsoi et al., 2014). Improving our understanding of how new neuron survival is regulated across brain regions and between hemispheres may inform strategies that consider interconnected effects in a whole brain system in enhancing the survival of new neurons post brain injury or disease (Mu and Gage, 2011; Sun, 2014; Winner and Winkler, 2015) and in promoting brain health more generally.

## Materials and Methods

### Animals

All methods were approved by the Queens College Institutional Animal Care and Use Committee (IACUC). Subjects were adult male zebra finches between 4 and 11 months of age. In zebra finches, only the males sing. We examined both the recruitment and survival of new neurons using doublecortin (DCX) and bromodeoxyuridine (BrdU) labeling, respectively. Different cohorts of birds were used in labeling either DCX and BrdU, with overlapping individuals used within a label: DCX in HVC (n = 20), Area X (n = 24), and NCM (n = 19); BrdU in HVC (n = 22), Area X (n = 40), and NCM (n = 22). Birds were hatched in either the Queens College or Wesleyan University breeding colony and kept with their parents until 90 days of age. Thereafter, birds were group housed, with experimental and control birds housed together in group cages, within auditory and visual contact of both sexes throughout the study. Birds were maintained on a 12:12 h light:dark schedule with food and water available *ad libitum*.

### Song Recording and Analysis

Adult male zebra finches were temporarily housed individually in sound attenuated chambers to record songs pre- and post-surgery. They were recorded continuously for 2-4 days prior to surgery, and again 1-2 days after surgery and then returned to their home cages. Recordings were made (Earthworks precision audio SR20 Cardioid Microphone) using sound-activated Avisoft Recorder (Avisoft Bioacoustics). Song bouts were edited to single motifs with Raven sound analysis software (Cornell Lab of Ornithology). Changes in structure after surgery were computed using Sound Analysis Pro (Tchernichovski et al., 2000). We used two measures to assess song quality: (1) accuracy and (2) percentage of similarity (termed here “similarity”). The accuracy score indicates the fidelity with which song elements were reproduced by comparing 1 ms song segments between pre- and post-operative motifs. Similarity indicates sameness in song structure at longer intervals (~50 ms) (http://soundanalysispro.com/manual-1). We compared 10 pre-surgery and post-surgery motif pairs for each bird and used the mean of these scores as a measure of postoperative song change.

We were interested in whether the degree of global song change post TS-cut, or a change in a particular song feature, corresponded to numbers of new neurons in various brain regions. Therefore, we ran independent linear regressions between the following measures of pre- to post-surgery song change and cell counts: accuracy, similarity, pitch difference, FM difference, entropy difference, and goodness difference. P values were corrected for multiple comparisons by adjusting the significance p value to 0.05/6 = 0.0083, using the Bonferroni correction. Uncorrected p values are reported in the text along with the corrected p value threshold necessary for significance.

Singing rates were computed manually from continual voice-activated recordings taken on days 2-4 post-surgery. Total songs were counted for each bird during a single hour within two hours of lights coming on. Mean songs per hour were calculated.

### Bromodeoxyuridine (BrdU) Injections and Tracheosyringeal Denervation

All birds received intramuscular injections of 5-Bromo-2’-deoxyuridine (BrdU; 74 μg/g, pH 7.4, Sigma) 3x/day for three days to label mitotically active cells. Surgeries were performed 21-22 days after the last BrdU injection, such that new neurons were 21-24 days old when song feedback was altered. This time frame was selected in order to ensure new neurons had migrated into the three regions of interest prior to nerve resection.

Birds were anesthetized with either a mixture of ketamine and xylazine (0.03-0.05 mg/g and 0.06 mg/g, respectively), or isoflurane in compressed air (3%; Henry Schein). Using a surgical microscope (Zeiss Universal S3B), a ~3-mm rostro-caudal incision was made < 1 mm lateral to the ventral midline to expose the tracheosyringeal nerve. A 2-4 mm section of either the left or right tracheosyringeal nerve (nXIIts, “TS”) was resected to prevent regrowth following surgery. Sham birds received all surgical manipulations except nerve resection.

### Histology

Birds experienced altered song for seven days and then were overdosed with sodium pentobarbital (Euthasol) and transcardially perfused with 0.1 M phosphate buffered saline (PBS, pH 7.4) followed by 4% paraformaldehyde (PFA; Sigma-Aldrich; pH 7.4). Brains were post-fixed for 1 h in 4% PFA, rinsed in PBS for 3 h and embedded in polyethylene glycol (MW = 1500; Polysciences). Six-ųm sagittal sections were cut on a rotary microtome. The brain was oriented with the midline parallel to the blade edge. The first complete section through the telencephalon was saved and subsequently every sixth section of tissue was mounted onto Superfrost Plus + slides (VWR). All series were stored at −20⁰ C prior to processing using immunohistochemistry (IHC).

### DCX Immunohistochemistry

Immunolabeling was performed using an anti-DCX goat polyclonal IgG (Santa Cruz Biotechnology; sc-8066, 1:150) or anti-DCX rabbit polyclonal IgG (Abcam; ab18723, 1:1000). Sections were brought to room temperature in TBS, followed by a 10-min wash in fresh TBS. Sections were then incubated for 30-min in a H_2_O_2_ solution (97% TBS, 1% methanol, 2% of 3% retail H_2_O_2_) to eliminate endogenous peroxidases. After three 5-min TBS rinses, sections were blocked with 5% normal horse serum (Jackson ImmunoResearch Laboratories Inc.) and 0.5% Triton X-100 (Sigma-Aldrich) in TBS for 30-min at room temperature, followed by exposure to anti-DCX polyclonal antibody in the same blocking buffer overnight. After three 5-min TBS rinses, sections were incubated for 3 h in biotinylated horse anti-goat (1:200, Vector Labs) or biotinylated goat anti-rabbit secondary antibody (1:200, Vector Labs) in TBS, rinsed again, and exposed for 1 h to an avidin-biotin complex (Vector Labs). Sections were rinsed in TBS and then reacted in a solution of 0.04% of 3,3’-diaminobenzidine tetrahydrochloride (DAB, Vector Labs) with nickel until the tissue changed color. Following three 5-min TBS rinses, sections were dehydrated in ethanols, delipidized in xylenes, and cover slipped with Krystalon (Millipore-Sigma).

### BrdU/Hu Immunohistochemistry

Sections were brought to room temperature in 0.1 M phosphate buffer (PB; pH 7.4), then exposed to citrate buffer (pH 5.6 – 6.0) at 90-95°C for 10 min, followed by a 5-min PB wash (37°C), 3 min in a solution of 0.28 % pepsin in 400 ml 0.1 M HCl at 37°C, and three 5-min washes in PB at room temperature. Sections were blocked with 3% normal donkey serum (Jackson Labs) and 0.5 % Triton X-100 in PB for 1 h at room temperature, followed by 24-48 h exposure to sheep anti-BrdU (1:239, Capralogics) in block at 4°C. Sections were again rinsed in PB and then incubated overnight in biotinylated donkey anti-sheep IgG in PB (1:200, Vector Labs), followed by overnight incubation in streptavidin-conjugated Alexa 488 in PB (1:800, ThermoFisher Scientific) for visualization of BrdU. The next day, sections were washed with PB, blocked for one hour, and incubated overnight in mouse anti-Hu primary antibody at 4°C (1:200 in 3% block; Invitrogen). After three 5-min PB rinses at room temperature, tissue was exposed to donkey anti-mouse IgG conjugated to Cy-3 in PB (1:80; EMD Millipore) for 1 h for visualization of Hu. Finally, sections were washed, dehydrated with ethanols, delipidized with xylenes, and cover slipped with Krystalon (Millipore-Sigma).

### Microscopy

Data were collected without knowledge of bird identity, treatment, or brain hemisphere. We had no predictions for potential hemispheric differences in new neurons in HVC or Area X. Area measurements and cell counts for all regions were performed using a computer-yoked microscope and mapping software (Olympus BX51; Lucivid LED microprojection, Neurolucida, MicroBrightField, Inc.). The boundaries of all brain regions were traced in 6-12 sections per hemisphere per bird. Boundaries for HVC were established with dark-field optics based on neuropil density and contrast. Area X is found ventral and anterior to the densely reflective oval shaped lateral magnocellular nucleus of anterior nidopallium (lMAN) and is demarcated by a dense haze of terminals and small cells. Area X and lMAN are separated by lamina pallio-subpallialis (LPS). NCM was identified as in (Pytte et al., 2010). DCX^+^ cells were visualized with bright field light microscopy (Figure 2A,B). BrdU^+^/Hu^+^ cells were visualized using fluorescein isothiocyanate (FITC) and rhodamine filters and a dual FITC/rhodamine filter (Figure 2C,D,E). New neurons per square millimeter was calculated by dividing the numbers of labeled cells by the total area sampled.

**Figure 2.**
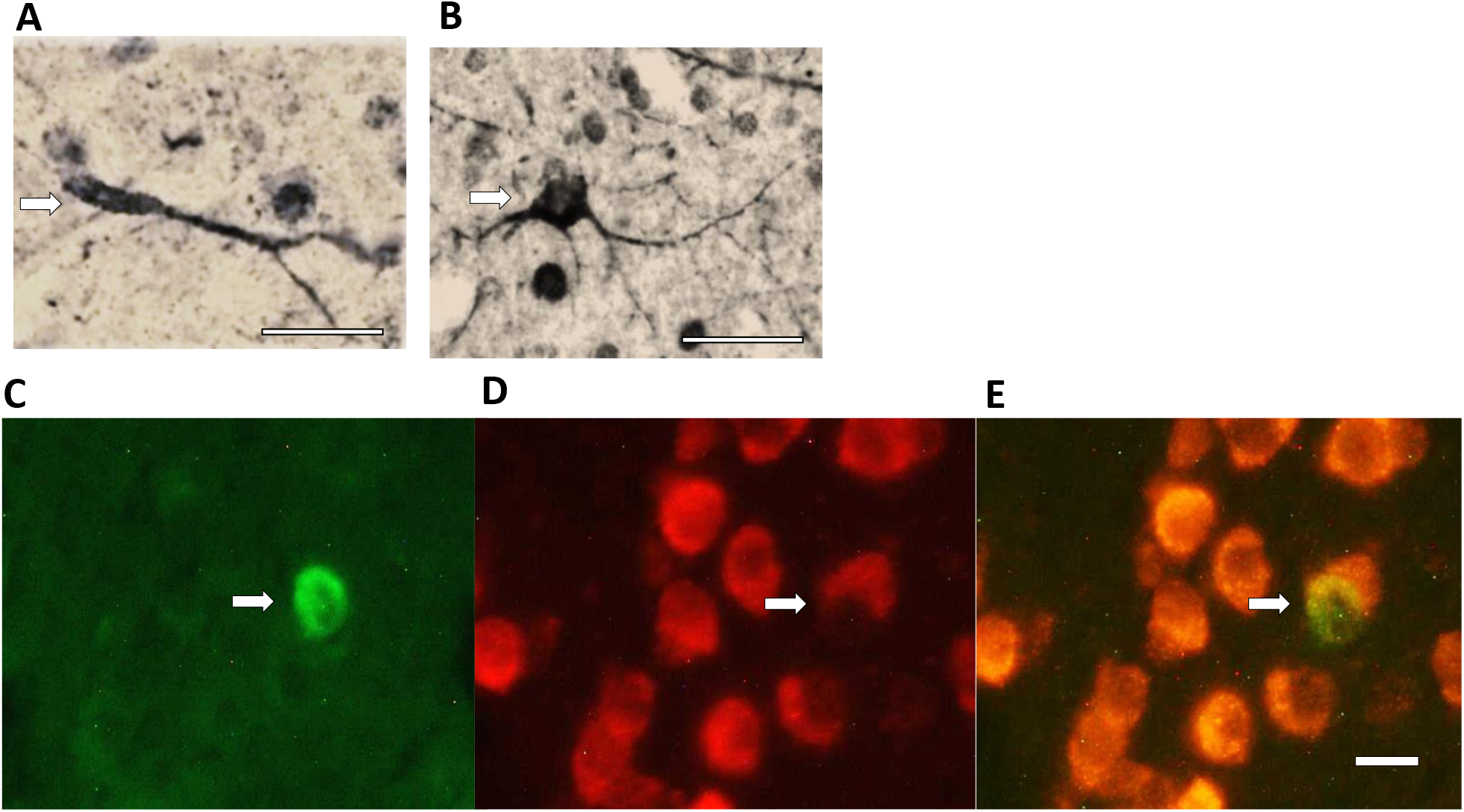
Photomicrographs of doublecortin-positive neurons in HVC. A. Fusiform, approximately 1-2 weeks old as indicated by its thin, elongated soma and unipolarity and B. Multipolar, approximately 2-3 weeks old as indicated by large, round soma and multiple processes (age estimates following Balthazart, et al., 2008). (C) Fluorescent markers used to identify 28-30 d old neurons in HVC, BrdU^+^ nucleus visualized with FITC filter (D) Hu^+^ neurons seen under rhodamine filter (E) BrdU^+^/Hu^+^ neuron showing colocalization of the two markers (rhodamine/FITC filter) in the same field of view. Scale bars A, B = 25 μm; C, D = 10 μm.

### Statistical Analysis

Data are presented as means and SEMs unless otherwise specified. Analyses were performed using one-way ANOVAs, two-factor ANOVAs with repeated measures on hemisphere, and two-tailed *t*-tests for independent samples. Tukey’s HSD post-hoc tests were used following significant effects determined by ANOVA. Following significant mixed ANOVAs, we performed one-way ANOVAs between groups within individual hemispheres and then Holm’s procedure with adjusted p values for multiple tests. Pearson correlations were computed between cell counts across brain regions or hemispheres. The lateralization index (LI) indicates the relative number of new neurons between hemispheres, normalized for the bird’s mean number of new neurons per area sampled in both hemispheres (as in Tsoi et al., 2014). Higher positive values indicate more new neurons in the left hemisphere relative to the right:

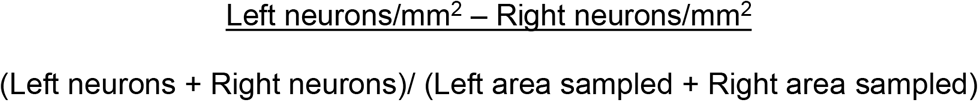

## Results

### Song Analysis

Birds that underwent either left or right TS-cut had significantly lower accuracy scores, a measure of song change post-denervation, than controls (One-way ANOVA, df = 2, F = 13.95, p = 0.002, Figure 3A). Both left TS-cut and right TS-cut scores were lower than those of controls (Tukey’s post-Hoc test, p < .01) and there was no difference between the two denervated groups (Tukey’s post-Hoc test p > .05). On the other hand, there was not a significant difference in pre-postop similarity scores among groups (one way ANOVA, df = 2, F = 1.84, p = 0.22, Figure 3B), suggesting that overall song structure was preserved in TS-cut birds despite degradation of fine structure. Post-operative singing rates did not differ among groups (One-way ANOVA, df = 2, F = 0.231, p = 0.797 data not shown).

**Figure 3.**
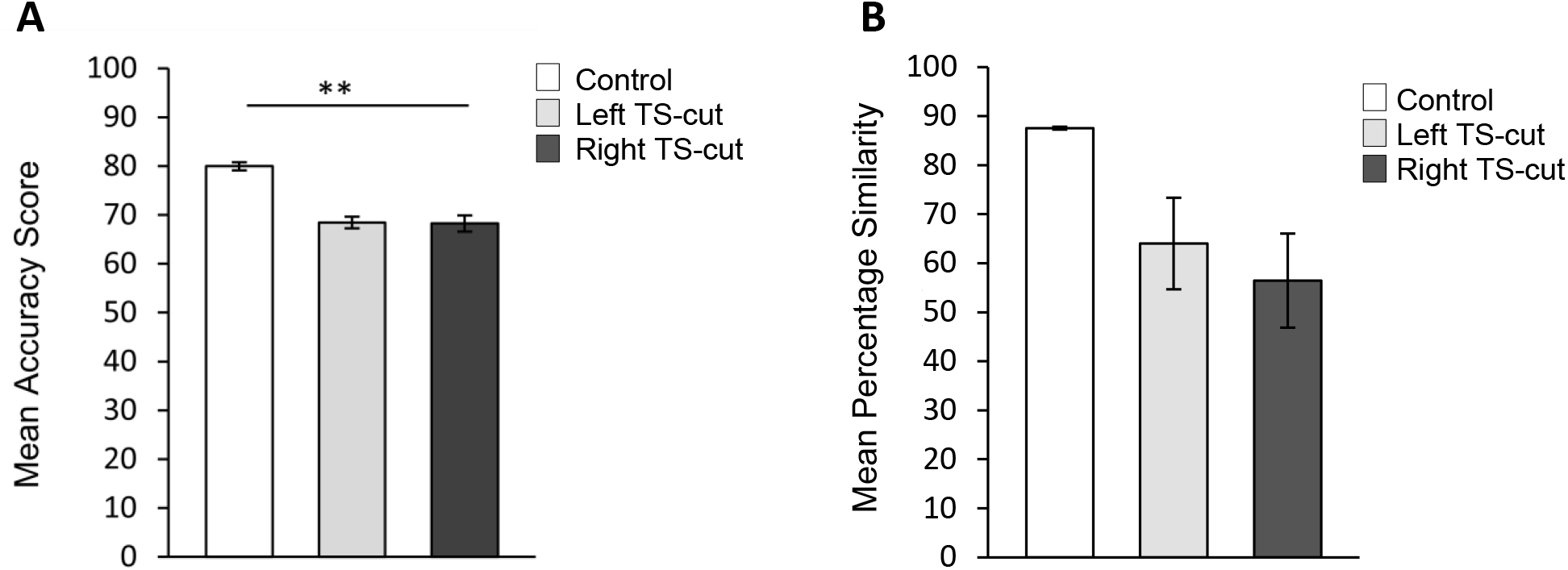
A. Effects of TS-cuts on song structure. A. Birds that underwent left (n = 5) or right (n = 4) TS-cut had accuracy scores that were significantly lower than those of controls (n = 2), demonstrating that unilateral tracheosyringeal nerve cut impacts the quality of song production. B. There were no significant differences in percentage similarity scores between pre-operative and post-operative songs of control (n = 2), left TS-cut (n = 5), and right TS-cut birds (n = 4). ** = p < .01

### Effect of TS-cut on New Neurons in HVC

There was no effect of treatment (df = 2, F = 0.75, p = 0.49), hemisphere (F = 0.04, p = 0.84), or interaction (F = 0.29, p = 0.75) on numbers of BrdU^+^/Hu^+^ cells in HVC that were 28-30 days old (two-factor ANOVA, repeated measures on hemisphere, Figure 4A). We also quantified DCX+ cells, a marker for immature neurons that is expressed for 2-3 weeks after mitosis in mammals (Brown et al., 2003) and was calculated to be expressed in newborn neurons for a similar time, approximately 20 days, in the canary (Balthazart, et al., 2008). The morphology of DCX+ neurons can also be used to estimate neuronal age in this time frame with fusiform, bipolar cells presumably with a migratory morphology being younger, and round, multipolar cells being older (Figure 2A, B; Balthazart, et al., 2008). We did not find treatment or hemisphere differences when fusiform and multipolar cell counts were analyzed independently (p > 0.05 for all); therefore, we combined DCX+ cells. We found no effect of treatment (df = 2, F = 0.16, p = 0.85), hemisphere (df = 1, F = 0.05, p = 0.83), or treatment by hemisphere interaction (df = 2, F = 0.39, p = 0.68) on numbers of DCX+ cells in HVC (two-factor ANOVA, repeated measures on hemisphere, Figure 4B). We also found no difference in new neuron numbers (neither BrdU+/Hu+, nor DCX+) between hemispheres ipsilateral and contralateral to the TS nerve cut.

**Figure 4.**
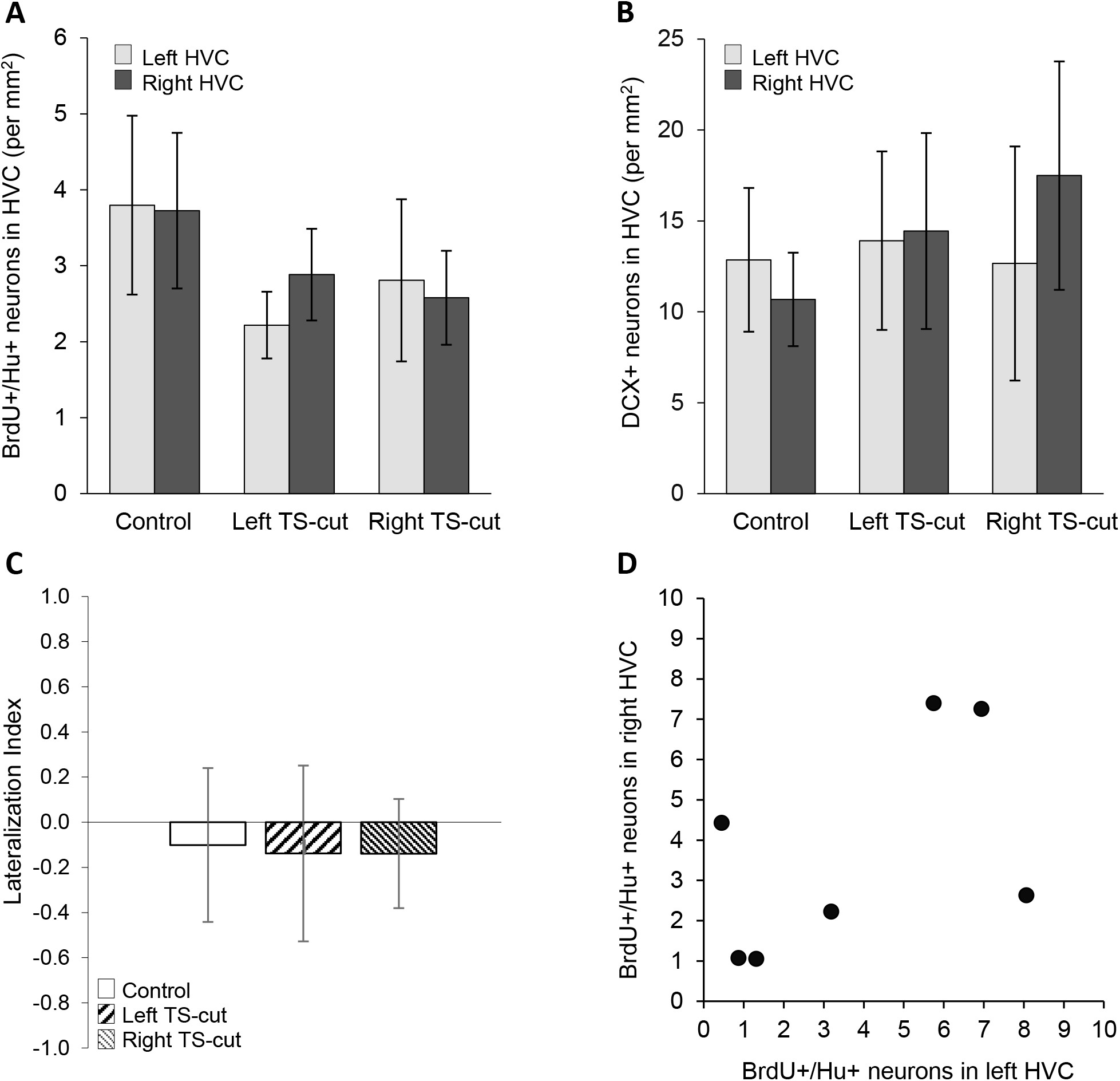
Numbers of new neurons in HVC as a function of hemisphere and experimental condition. A. Neither left (n = 7) nor right (n = 8) TS-cut affected the number of 28-30 d old BrdU+/Hu+ neurons in HVC compared to controls (n = 7). B. Neither left (n = 8) nor right (n = 5) TS-cut affected the number of ~1-3 week old DCX+ neurons in HVC compared with controls (n = 7). C. There were no differences in the Lateralization Index of BrdU+/Hu+ cells among treatment groups. D. There was no correlation between numbers of BrdU+/Hu+ cells between the left and right HVC.

Unilateral TS-cuts had no effect on the lateralization index of BrdU+/Hu+ or DCX+ neurons in HVC (one-way ANOVA, df = 2, F < 0.01, p = 1.0; df = 2, F = 1.19, p = 0.33, respectively, Figure 4C). We also found no relationship between the number BrdU+/Hu+ neurons or DCX+ cells in left HVC and right HVC of control animals (df = 5, r = 0.53, t = 1.38, p = 0.23; df = 5, r = 0.66, t = 1.1, p = 0.32, respectively, Figure 4D).

### Effect of TS-cut on New Neurons in NCM

There was no main effect of treatment (F = 0.22, p = 0.80), hemisphere (F = 1.29, p = 0.27), or 3 way treatment by hemisphere interaction (F = 0.83, p = 0.45) on numbers of BrdU+/Hu+ neurons in NCM (Figure 5A). As in HVC, we found no difference among treatments or hemispheres when DCX+ fusiform and multipolar cells were analyzed separately; therefore, we combined these categories. There was no main effect of treatment (F = 0.48, p = 0.63), hemisphere (F = 2.28, p = 0.15), or interaction on DCX+ cells in NCM (F = 0.14, p = 0.87, two-factor ANOVA, repeated measures on hemisphere, Figure 5B).

**Figure 5.**
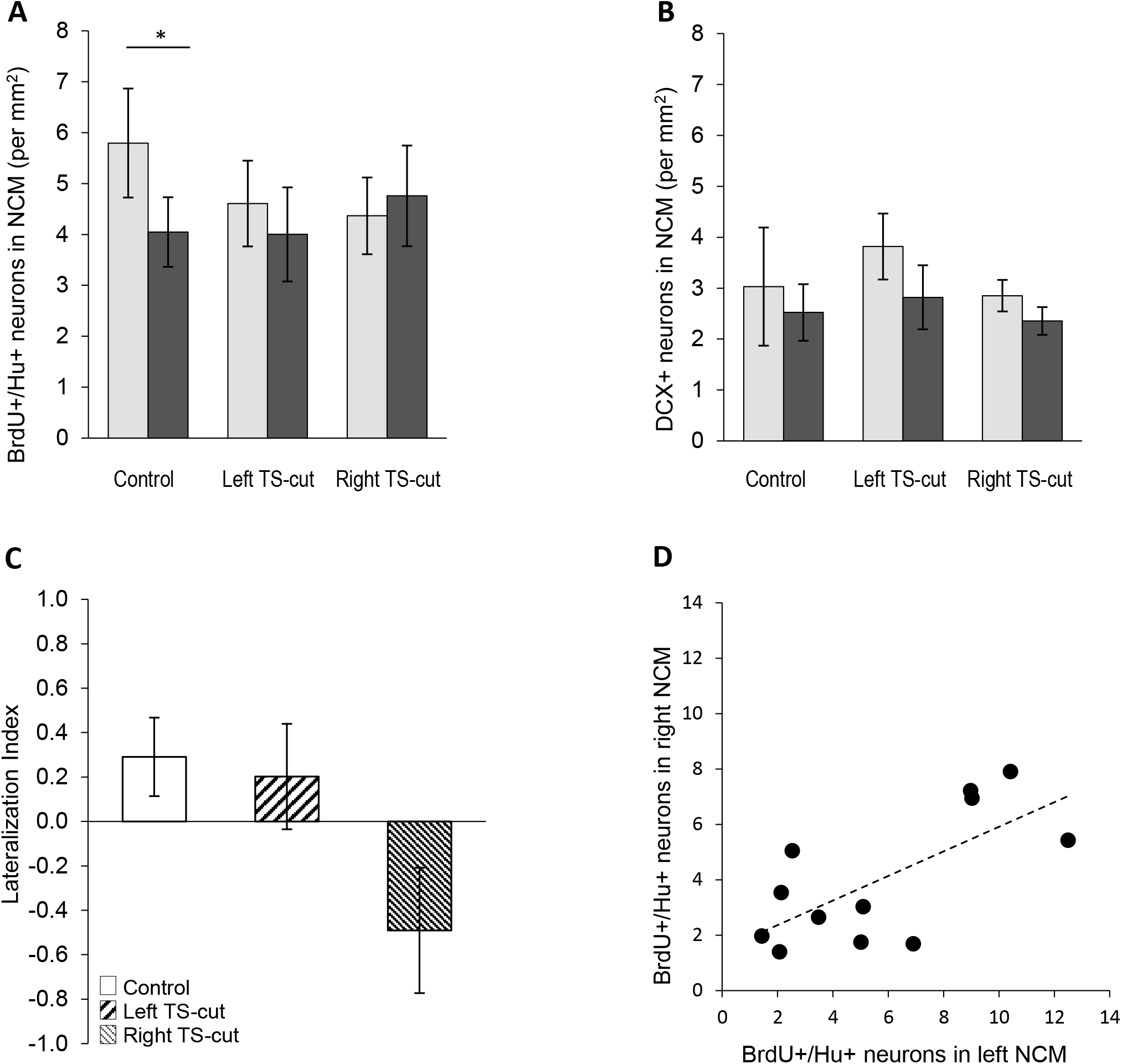
Numbers of new neurons in NCM as a function of hemisphere and experimental condition. A. Left (n = 12) and right (n = 11) TS-cuts had no effect on the overall number of 28-30 d old BrdU+/Hu+ neurons in NCM. Control birds (n = 12) had more BrdU+/Hu+ neurons in left NCM than the right and this asymmetry was not seen in the left or right TS-cut group. B. Neither left (n = 7) nor right (n = 6) TS-cut differed from controls (n = 7) in numbers of DCX+ neurons in NCM. C. There was a positive correlation between numbers of 28-30 d old neurons in left NCM and right NCM of control birds. * = p < 0.01.

Earlier work showed that in unmanipulated controls, there were more new neurons in the left NCM than the right, and that this asymmetry was lost after a unilateral tracheosyringeal nerve cut (Tsoi et al., 2014). Likewise, we found more BrdU^+^/Hu^+^ neurons in the left than the right NCM (paired sample t-test, t (11) = 2.26, p = 0.045) in controls. Hemispheric asymmetry was not seen in either the left TS-cut or right TS-cut group (paired t-test within group, p = 0.67, p = 0.74, respectively). Consistent with this pattern, we found a significant difference in lateralization indices of BrdU+/Hu+ among the three groups (df = 2, F = 3.92, p = 0.03). Interestingly, the lateralization index of only the right TS-cut group differed from controls (HSD, p <0.05) whereas the lateralization index of the left TS-cut group did not differ from that of controls or the right TS-cut group (Figure 5C).

Unlike in HVC, in control birds, the numbers of BrdU+/Hu+ cells in left and right NCM were positively correlated (df = 2, r = 0.69, t = 3.06, p = 0.01, Figure 5D) as in Tsoi et al. (2014). There were no correlations between NCM hemispheres in either TS-cut group (p = 0.45, left TS cut; p = 0.63, right TS-cut, data not shown). There were no correlations in numbers of DCX+ cells between NCM hemispheres in any group (p > 0.05 for all, data not shown).

### Effect of TS-cut on New Neurons in Area X

We found a significant difference in the number of BrdU+/Hu+ neurons in Area X among the three treatment groups (two factor repeated measures ANOVA, F = 7.47, p = 0.002, Figure 6A). There was no difference between hemispheres (F = 0.25, p = 0.62) and no interaction (F = 1.16, p = 0.33). Within the left hemisphere (one way ANOVA, df = 2, F = 3.31, p = 0.048), numbers of BrdU+/Hu+ cells in both the left TS-cut and right TS-cut decreased compared to controls (t = 2.05, p = 0.048; t = 2.39, p = 0.045, respectively). The same pattern was found within the right hemisphere Area X (one way ANOVA, df = 2, F = 6.84, p = 0.003). Compared to the right Area X of controls, there were fewer numbers of BrdU+/Hu+ cells in the right Area X of both the left TS-cut and right TS-cut groups (t = 3.58, p = 0.002; t = 2.65, p = 0.012, respectively).

**Figure 6.**
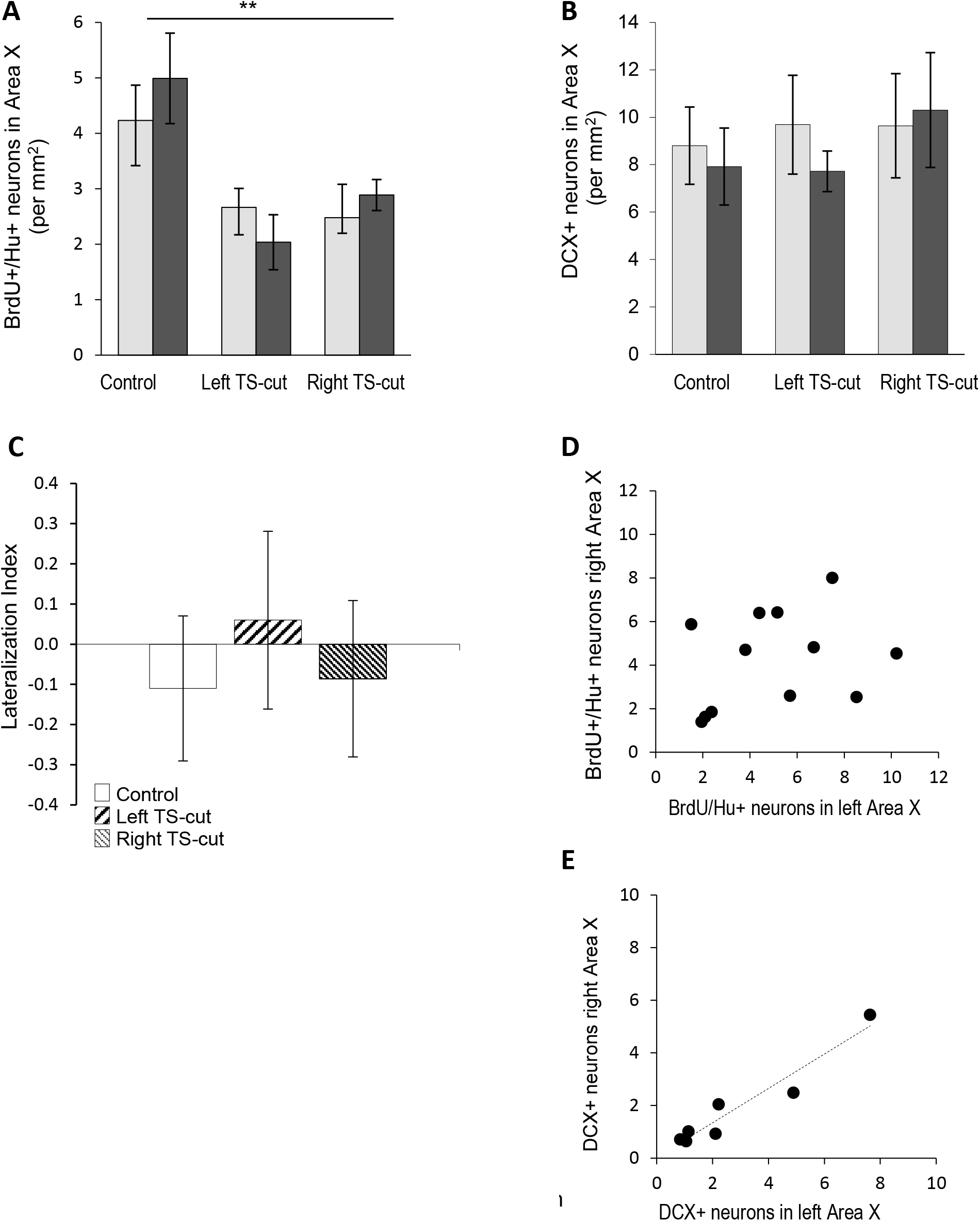
Numbers of new neurons in Area X as a function of hemisphere and experimental condition. A. Left (n = 12) and right (n = 14) TS-cuts significantly decreased the number of 28-30 day old neurons compared to controls (n = 12). B. Neither left (n = 9) nor right (n = 8) TS-cut groups differed from controls (n = 7) in numbers of DCX+ neurons in Area X. C. Lateralization indices for BrdU+/Hu+ neurons in Area X. LIs did not differ between controls, left TS-cut, and right TS-cut conditions. D. There was no relationship between the numbers of BrdU+/Hu+ neurons in left versus right Area X in controls. E. We found a significant correlation between left and right Area X in numbers of DCX+ neurons in controls. ** = p < .01

As in the other regions, we combined all DCX+ cells after finding no difference within fusiform and round DCX+ cell types across treatments. There was no main effect of treatment (2 factor mixed ANOVA, F = 0.07, p = 0.93), hemisphere (F = 1.48, p = 0.24), or interaction (F = 1.22, p = 0.32; Fig. 6B) on the number of DCX+ neurons in Area X.

There was no significant difference among groups in the LI of BrdU+/Hu+ or DCX+ cells (one-way ANOVA, F = 0.2, p = 0.82; F = 0.20, p = 0.82, respectively, Figure 6C). There were no correlations between the numbers of BrdU+/Hu+ neurons in the left and right Area X of control animals (df = 10, r = 0.30, p = 0.35; Fig. 6D). However, DCX+ cells were correlated between hemispheres in control birds (df = 5, r = 0.92, p = 0.016, Figure 6E) and in birds with a left side TS-cut (df = 7, r^2^ = 0.85, p = 0.03), but not in birds with a right side TS-cut (df = 6, r = 0.67, p = 0.27, data not shown).

### Within Hemisphere Correlations in Numbers of New Neurons Between Adjacent Regions

We were also interested in whether new neuron numbers may be linked between functionally associated regions in control birds. However, we found no correlations in numbers of BrdU+/Hu+ cells between NCM and HVC or between HVC and Area X in either the left (df = 4, r = 0.69, p = 0.34; df = 4, r = 0.89, p = 0.063, respectively) or right (df = 4, r = 0.69, p = 0.35; df = 5, r = 0.49, p = 0.61, respectively) hemispheres. It is worth noting, however, that in each hemisphere removing a single bird would have resulted in significant correlations between HVC and Area X and thus larger sample sizes may find this to be the case. There were no correlations between DCX+ cells within either hemisphere between NCM and HVC (left: df = 4, r = 0.48, p = 0.34; right: df = 4, r = 0.47, p = 0.35) or between HVC and Area X (left: df = 4, r = 0.79, p = 0.063; right: df = 5, r = 0.23, p = 0.612, data not shown).

## Discussion

This is a broad-strokes study, serving as a first pass in understanding whether the quality of song structure affects new neuron survival and whether regulation of new neuron survival is independent across brain region and hemisphere. We found that irreversible disruption to song structure by unilateral denervation of the syrinx in adult male zebra finches impacted the numbers of 28-30 day old neurons in a region and hemisphere-specific manner. Altered song feedback had no effect on numbers of one month old neurons in HVC, resulted in a loss of left-sided dominance of new neurons in NCM consistent with Tsoi et al. (2014), and decreased neuron survival in both hemispheres of Area X. These effects were not present in younger neurons expressing DCX perhaps due to the lack of functional connectivity in this cohort at the time of treatment.

### HVC

The clearest example of a behavior that impacts new neurons in HVC is singing (Li et al., 2000; Alvarez-Borda and Nottebohm, 2002; Ball et al., 2002; Alward et al., 2016). However, functional implications of upregulating new neurons during singing are not clear. In the non-seasonal zebra finch, new neurons appear to be added in surplus of replacing dying cells as HVC neuronal packing density increases with increasing bird age (Walton et al., 2012). Therefore, song production, and the increased density of adult-formed neurons, may have the outcome of fine tuning motor stereotypy, or improved resolution of fine motor gestures, increasing the precision of song structure c.f.(Polomova et al., 2019). Consistent with the idea that new neurons contribute to precision, song stereotypy continues to increase with age in adult zebra finches (Brainard and Doupe, 2001; Pytte et al., 2007; James and Sakata, 2019). The proposal that new neurons in the HVC-RA motor pathway increase the motor resolution for song production borrows from the model that new neurons in the rodent hippocampus contribute to memory resolution, providing improved pattern separation (Aimone et al., 2011) and suggests a potential general rule of new neuron function in widely different substrates.

With increased bird age, and thus increased packing density of HVC-RA neurons (Walton et al., 2012), the motor pattern for song production is also increasingly resistant to degradation after deafening (Lombardino and Nottebohm, 2000). Lombardino and Nottebohm (2000) speculated that each time a bird sings, learning, or “motor engrainment”, cumulatively strengthens the memory of the song motor commands, decreasing online reliance on auditory feedback. Combining these findings, perhaps the act of singing, by upregulating new neurons, increases the substrate density for encoding the motor pattern, resulting in greater motor pattern precision and durability. This is consistent with the finding that more new neurons in HVC correspond to a more stable song post-deafening (Pytte et al., 2012).

Independent of this model but compatible with it, it has been suggested that as songs progress toward a target goal during development, seasonally, or with increased stereotypy with age, feedback from continually improving song quality may actively promote new neuron survival in HVC (Wilbrecht and Kirn, 2004; Kirn, 2010). Here we examined whether altered song feedback, without song improvement, affected numbers of new neurons. We found there was no group effect of the TS-cut on numbers of new neurons in HVC, examining the left or right hemisphere independently and combined, and also separately assessing the HVC hemisphere ipsilateral and contralateral to the TS nerve cut. Thus the negative results reported here are consistent with the alternative hypothesis that song improvement, i.e., progress toward a target, rather than altered feedback per se is necessary for increased survival of new neurons in HVC (Pytte et al., 2011).

### NCM

NCM is thought to play a role in the recognition of conspecific songs (Chew et al., 1996) and to encode at least one of potentially several distributed copies of a tutor’s song used in song learning (Solis and Doupe, 1997; Phan et al., 2006; Bolhuis and Moorman, 2015). Neurons in NCM selectively respond to conspecific songs, the song of the bird’s tutor, as well as playback of the bird’s own song (Grace et al., 2003; Hsu et al., 2004). Moreover, NCM is active while a bird is singing, suggesting it may process auditory feedback from song in real time (Jarvis et al., 1998). Motor recovery from song distortion requires an intact NCM, although it is not clear whether this reflects access to a stored representation of the bird’s own song or that of the tutor (Canopoli et al., 2014).

NCM receives a robust influx of new neurons throughout adulthood, although very little is known about the function of new neurons added. We know that new neurons are upregulated in birds housed in groups (Lipkind et al., 2002; Barnea et al., 2006; Adar et al., 2008a; Adar et al., 2008b) and are diminished in birds that have been deafened (Pytte et al., 2010). Together, this suggests that activity in NCM promotes new neuron survival; however, does not shed light on function. In a careful study of new neuron survival across the rostral-caudal extent of NCM, Barnea et al. (2006) proposed an interesting model describing how new neurons may encode information about the calendar timepoint when an event (song stimulus) was experienced. More generally, the continual addition of new neurons may preserve and/or improve the resolution of stored song memories in either hemisphere of NCM, and perhaps enhance song discrimination in general.

Our interest is in whether new neuron survival may be influenced by the degree to which song feedback matches expected feedback, perhaps with respect to a stored template of the bird’s own song. In control birds, we found more new neurons in the left than right NCM, consistent with Tsoi et al., (2014). This left-side lateralization in new neurons was lost in the TS-cut bird. Although not statistically significant, the loss in lateralization was due more to a decrease in numbers of new neurons in the left NCM than an increase in new neurons in the right NCM. Thus, we hypothesize that a template of the bird’s own song may be lateralized to the left, rather than right, NCM.

Evidence for a lateralized response in NCM to the bird’s own song is mixed. Interestingly, estradiol synthesis in the left NCM but not the right NCM is necessary for a bird’s preference for his own song (Remage-Healey et al., 2010). In a behavioral study, adult male zebra finches showed deficits in discriminating their own song from that of a cage-mate when the left side ovoidalis, part of the ascending auditory pathway, was lesioned and not when the right side was lesioned (Cynx et al., 1992). However, selectivity for playback of the bird’s own song was found to be lateralized toward the right side in the midbrain dorsal lateral mesencephalic nucleus (MLd) the analog of the mammalian inferior colliculus (Poirier et al., 2009). Like ovoidalis, MLd is part of the unilateral ascending auditory pathway that eventually projects to NCM. Moreover, both left and right NCM respond to playbacks of the bird’s song (Soyman and Vicario, 2017).

In a different model, hemispheric asymmetry may be based on sound acoustic features rather than categories of own song vs those of others (Van Ruijssevelt et al., 2017). Playback of conspecific songs with filtered spectral structure increased BOLD responses in the left NCM and slightly decreased BOLD responses in the right NCM compared to nonmanipulated songs (Van Ruijssevelt et al., 2017). Because spectral filtering removed frequency information from the song leaving primarily temporal content, the authors interpreted this finding as indicating that the left NCM is specialized for processing temporal information. They suggested that this asymmetry is normally masked by the spectral content of whole songs (Van Ruijssevelt et al., 2017). On the other hand, we suggest that perhaps the increased response in the left NCM reflects a left-dominant sensitivity to aberrant information, resonating with our findings. The left hemisphere of NCM has been shown to be sensitive to novel auditory experience more broadly (Moorman et al., 2015; Yang and Vicario, 2015) consistent with the idea that NCM (bilaterally) is sensitive to expectation (Lu and Vicario, 2017; Dong and Vicario, 2018). Thus perhaps it is the novelty of the distorted song, and not feedback about the bird’s own song *per se,* that underlies the observed decrease in new neurons in the left NCM.

It is a consistent finding across studies that the magnitude of asymmetry in various measures of NCM corresponds to performance in learning (Moorman and Nicol, 2015). For instance, the degree of left side dominance in new neurons (Tsoi et al., 2014), and left side dominance in activity during sleep (Moorman et al., 2015) predict accuracy of song learning of the bird’s own song. Left side dominance in NCM activity in response to song playback is also correlated with rate of learning in a conspecific song discrimination task (Bell et al., 2015). The results of the current study and that of Tsoi et al., (2014) add that lateralization in new neurons is flexible and can be disrupted by the bird’s experience.

### AREA X

We found significantly fewer new neurons in both hemispheres of Area X in birds who received TS nerve cuts. Area X plays a role in juvenile song learning (Sohrabji et al., 1990; Scharff and Nottebohm, 1991; Olveczky et al., 2005) and modification of the song motor pattern in adults in response to social context, deafening, and perturbations in feedback, although the mechanism by which it does so is debated (Hessler and Doupe, 1999; Brainard and Doupe, 2000; Ali et al., 2013; Woolley et al., 2014; Woolley and Kao, 2015; Woolley, 2016; Kojima et al., 2018) c.f. (Sanchez-Valpuesta et al., 2019). In adult birds, the anterior forebrain pathway (AFP) more generally mediates differences in song variability (Hessler and Doupe, 1999; Williams and Mehta, 1999; Woolley et al., 2014).

As in HVC (Walton et al., 2012) new neurons are continually added to Area X, increasing packing density with age, and are incorporated into functioning circuits (Kosubek-Langer et al., 2017). Approximately 80% of new neurons express D1 and D2 receptors, become medium spiny neurons (MSN), and fire during singing (Kosubek-Langer et al., 2017). New MSN are also innervated by glutamatergic synapses, putatively from HVC and lMAN, by day 31 after birth dating (Kosubek-Langer et al., 2017). Kosubek-Langer et al., (2017) suggest that the newly recruited MSNs fulfill the same function as the existing older MSNs. These authors speculate that the continual addition of MSNs results in relatively stronger inhibition by this increasing population on the unchanged number of pallidal-like neurons. Downstream DLM then would diminish excitation of LMAN and subsequently, of RA. Kosubek-Langer et al., (2017) note that this model predicts diminishing AFP influence over the motor pathway as birds age, consistent with behavioral studies (Lombardino and Nottebohm, 2000; Brainard and Doupe, 2001; Pytte et al., 2007).

Song variability or degradation of acoustic structure can be induced experimentally by distorting auditory feedback via tracheosyringeal nerve cut or deafening (Nordeen and Nordeen, 1992; Williams and McKibben, 1992; Hough and Volman, 2002; Nordeen and Nordeen, 2010). The AFP seems to mediate this degradation process, since lesions of the AFP output nucleus LMAN prevent song deterioration after auditory feedback distortion (Brainard and Doupe, 2000; Nordeen and Nordeen, 2010). In addition, our results suggest that addition and/or survival of new neurons in Area X is not immune to behavioral feedback and may be influenced by the quality of song structure.

The mechanism by which altered song may result in decreased survival of new neurons in Area X is not known, but may involve modulation of dopamine. Dopamine interacts with mechanisms of new neuron survival (Appeltants et al., 2000; Cornil et al., 2008; Leblois et al., 2010) and plays an important role in learning behaviors that are reinforced by rewarding outcomes. Most of the song system nuclei receive dopaminergic input (Lewis et al., 1981; Bottjer, 1993; Appeltants et al., 2000; Appeltants et al., 2002). In particular, the substantia nigra pars compacta (SNc) and the ventral tegmental area (VTA) send direct dopaminergic projections to Area X (Sasaki et al., 2006). Through evaluation of predicted and actual auditory feedback, perhaps dopaminergic signals modulate AFP output and influence song variability and learning.

Accurate song feedback may maintain dopamine activity and, in turn, new neuron survival. Activation of dopaminergic neurons in Area X occurs in several contexts (Scharff and Nottebohm, 1991; Kao and Brainard, 2006; Sasaki et al., 2006), including in response to playback of the bird’s own song (Gale and Perkel, 2010). It has previously been demonstrated that singing induces dopamine release from SNc and VTA into Area X (Sasaki et al., 2006). Likewise, it has been found that Area X may make an indirect connection to SNc/VTA via a projection to ventral pallium (VP) (Gale et al., 2008). Through this indirect pathway, the modulation of Area X output drives changes in the firing rate of dopaminergic neurons in the SNc and VTA via the VP. This relationship may suggest that Area X plays a role in feedback evaluation of song and transmits this information to dopaminergic neurons. Perhaps this pathway results in decreased dopamine signaling in the TS-cut birds with mismatched feedback and fewer new neurons in Area X. Midbrain dopaminergic neurons send profuse projections to the striatum, including Area X (Kitt and Brauth, 1986; Durstewitz et al., 1999), and much sparser projections to the pallium, which includes HVC and NCM (Kitt and Brauth, 1986). Therefore, dopamine may be less influential in mediating new neuron survival in HVC and NCM.

### Numbers of New Neurons Between Regions

Few studies have made inter-regional comparisons of adult neurogenesis in the mammalian brain. Notably, however, prenatal stress has been shown to decrease neurogenesis in the mouse hippocampus in adulthood, with no effect on neurogenesis in the olfactory bulb (Belnoue et al., 2013). In the aging pigeon, the extent of new neuron decline is greater in the olfactory bulb than in the hippocampus, again differentiating regulatory mechanisms between neurogenic regions (Meskenaite et al., 2016). Rates of decline in new neurons in young adults also differ between HVC and Area X (Pytte et al., 2007). Our results are consistent with the idea that regulatory mechanisms operate independently across brain regions, even those linked functionally or anatomically.

### Taken together

Song learning by juveniles requires incremental improvement toward a target motor behavior and requires both Area X and HVC, in concert with neuronal tuning to the tutor and the bird’s own song in NCM. Moreover, it occurs at a time when new neuron incorporation is high. In one of many models of song learning, akin to procedural motor learning more broadly, Area X and the AFP is thought to evaluate feedback and influence subsequent motor commands produced by HVC, and the output of feedback evaluation is thought to influence subsequent motor commands. Our findings suggest that after the juvenile stage of song learning, the accumulation of new neurons in Area X may correspond directly to the quality of song feedback in matching to the target own-song template. On the other hand, new neurons in HVC are not culled or maintained based on the output of a match between expected and received feedback. New neurons in HVC may instead either 1) be influenced via feedback in judging progress toward a goal of an ideal song or 2) are not affected by song feedback at all, and instead new neurons may drive progress toward a goal song in a feedforward direction only (Pytte et al., 2011). As in bilateral Area X, new neurons in left NCM may be culled when song feedback no longer matches expected feedback of the bird’s own song. Thus, perhaps increasing the resolution of the neuronal substrate in the song system and NCM by increasing new neurons serves to achieve increased song stereotypy (Pytte et al., 2007) both by better resolution of song feedback evaluation and/or by fine tuning motor output.

### Conclusions

This study contributes to our understanding of whether mismatched behavioral feedback affects new neuron recruitment and survival in brain regions that subserve and monitor those behaviors. We found that altering song production via unilateral tracheosyringeal denervation resulted in decreased 28-30 day old neurons in Area X, loss of lateralization of neurons in NCM, and had no effect in HVC. This indicates that the effects of syringeal denervation on neurogenesis vary depending both on the region and hemisphere that receives new neurons.

## Acknowledgements

John Kirn generously gifted birds for the study. Ben Koo contributed to tissue processing and cell quantification.

This article is dedicated to John R. Kirn (1952-2019).

